# Pf bacteriophages hinder sputum antibiotic diffusion via electrostatic binding

**DOI:** 10.1101/2024.03.10.584330

**Authors:** Qingquan Chen, Pam Cai, Tony Hong Wei Chang, Elizabeth Burgener, Michael J. Kratochvil, Aditi Gupta, Aviv Hargil, Patrick R. Secor, Josefine Eilsø Nielsen, Annelise E. Barron, Carlos Milla, Sarah C. Heilshorn, Andy Spakowitz, Paul L. Bollyky

## Abstract

Despite great progress in the field, chronic *Pseudomonas aeruginosa* (*Pa*) infections remain a major cause of morbidity and mortality in patients with cystic fibrosis, necessitating treatment with inhaled antibiotics. Pf phage is a filamentous bacteriophage produced by *Pa* that has been reported to act as a structural element in *Pa* biofilms. Pf presence has been associated with resistance to antibiotics and poor outcomes in cystic fibrosis, though the underlying mechanisms are unclear. Here, we have investigated how Pf phages and sputum biopolymers impede antibiotic diffusion using human sputum samples and fluorescent recovery after photobleaching. We demonstrate that tobramycin interacts with Pf phages and sputum polymers through electrostatic interactions. We also developed a set of mathematical models to analyze the complex observations. Our analysis suggests that Pf phages in sputum reduce the diffusion of charged antibiotics due to a greater binding constant associated with organized liquid crystalline structures formed between Pf phages and sputum polymers. This study provides insights into antibiotic tolerance mechanisms in chronic *Pa* infections and may offer potential strategies for novel therapeutic approaches.

**Teaser:** Pf phages and sputum polymers reduce antibiotic diffusion via electrostatic interactions and liquid crystal formation.

## Introduction

Cystic Fibrosis (CF) is a life-shortening disease marked by chronic pulmonary infections, inflammation, and structural damage of the lung tissue (*1*). The absence or dysfunction of the epithelial cystic fibrosis transmembrane conductance regulator (CFTR) protein leads to a buildup of viscous mucous in the airways (*2*). While CFTR modulator therapy has improved outcomes significantly for people with CF in many ways, infections remain problematic (*3*).

*Pseudomonas aeruginosa (Pa)* is the most common pathogen in bacterial lung infections by adulthood for people with CF (pwCF). Chronic endobronchial infection with *Pa* is associated with poor lung function, increased pulmonary exacerbations, and increased mortality (*4–7*). While *Pa* eradication protocols are now standard of care, success is unfortunately variable and not sustained (*8, 9*). Nearly 60% of patients will have chronic *Pa* infection by their mid-20s (*10*). Despite the advent of CFTR modulator therapy, chronic *Pa* infections remain an important problem in CF (*11*).

Tobramycin (TOB) is the most common inhalational antibiotic used to treat chronic *Pa* infections, though other anti-pseudomonal antibiotics are often used (*12*). However, clinical responses to antibiotics in pwCF are frequently variable and not necessarily predicted by antibiotics susceptibility testing (AST) data (*13–15*). Therefore, there is great interest in identifying factors that impact antibiotic efficacy in CF.

We and others have identified a role for Pf bacteriophages – a virus which infects *Pa* – in *Pa* pathogenesis and susceptibility to antibiotics. Pf phages are filamentous bacteriophages of the Inovirus family (*16, 17*) that are widespread across *Pa* isolates (*18–20*). Unlike most phages that parasitize bacteria (*21, 22*), filamentous phages like Pf are pseudolysogenic – they do not require the lysis of their bacterial hosts to release infectious virions (*23*). Instead, virions are extruded from the cell envelope of *Pa* without lysis. There are indications that Pf may enhance *Pa* fitness (*24, 25*) and promote *Pa* biofilm formation (*26*). *Pa* biofilms that include Pf phages contain liquid crystalline regions with alignment of the Pf filaments (*27, 28*) driven by depletion attraction forces between Pf phages and polymers (*25*). Pf mixed with polymers present during *Pa* infection, such as DNA or hyaluronic acid, rapidly assemble into thick, adherent, birefringent biofilms (*29*), indicating that the liquid crystalline state seen in *Pa* biofilms stem from interactions between Pf phages and polymers.

Pf phage is associated with more severe disease exacerbations in pwCF (*30, 31*) and chronic wound infections (*32*). Pf also reduces antibiotic efficacy (*33*) as Pf mixed with polymers typically found in human sputum increased *Pa* tolerance to tobramycin, gentamicin, and colistin (*27*). Moreover, Tarafder *et al.* showed visually how Pf phages form occlusive sheaths around *Pa* that shield them from antibiotics (*34*). Phage-mediated polymer organization may be particularly relevant to the pathophysiology of CF airway infections, where dense, polymer-rich secretions in the airways create ideal conditions for liquid crystal formation.

The mechanism underlying how Pf phages hinder antibiotic efficacy is unclear. Rossem *et al.* proposed a mathematical model wherein antibiotic diffusion through a single liquid crystalline tactoid, formed by Pf phages and polymers and encapsulating a bacterium, is limited by the adsorption of antibiotics to Pf phages (*35, 36*). While the general trend of their modeling agreed with previously published data on antibiotic and phage interactions, we sought to perform a more direct comparison between experimental and model results to illuminate the underlying mechanisms. Thus, in addition to our experimental observations, we also developed a set of continuum models for the entire sputum environment (encompassing tactoids that can form within) to hypothesis test potential mechanisms.

In this work, we have asked how Pf phages interact with commonly used inhaled antibiotics, such as tobramycin, in a novel model system incorporating sputum samples and sputum polymers. Using fluorescent recovery after photobleaching (FRAP), calorimetry, and mathematical modeling, we show that Pf inhibits antibiotic diffusion via charge-based interactions. These findings may help guide the selection of inhaled antibiotics and inform potential therapies directed at Pf phages.

## Results

### Tobramycin diffusion is impaired in Pf-positive sputum

We sought to define the impact of Pf phages on antibiotic diffusion in sputum. To this end, TOB was fluorescently labeled using Cy5 (Cy5-TOB) and homogenously mixed with sputum. We then used fluorescent recovery after photobleaching (FRAP), a technique where a laser is used to photo-bleach an area of the sputum mixture. After photo-bleaching, the fluorescence recovery of the bleached area is measured as a function of time as fluorescent particles diffuse from outside the bleached area into the bleached area (**Fig. 1A**). We obtained a fluorescence recovery curve from FRAP, which allows understanding of the diffusion rate of the fluorescently-tagged particle within the medium.

**Fig. 1.**
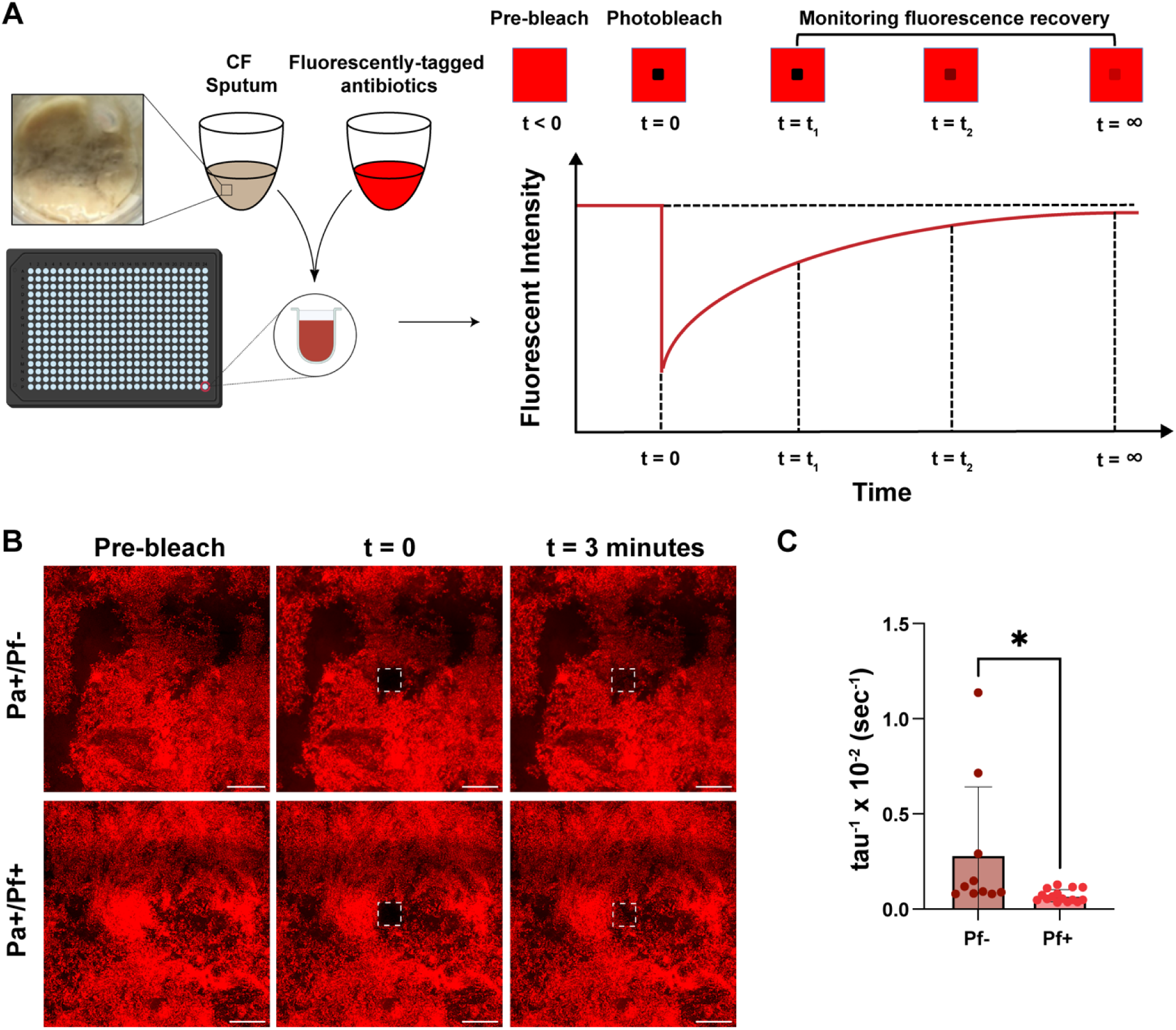
Tobramycin diffusion is decreased in Pf-positive sputum. **(A)** Fluorescent recovery after photobleaching (FRAP) was adopted to determine the diffusion of Cy5-tobramycin (Cy5-TOB) in sputum samples collected from cystic fibrosis patients. **(B)** FRAP was used to measure the recovery intensity of Cy5-TOB after photobleaching in sputum collected from Cystic Fibrosis (CF) patients. (**C)** The effective diffusion coefficient (tau^−1^) of Cy5-TOB in 3 patient sputum samples supplemented with Pf (Pf+, n=15) phages was less compared to Pf negative (Pf-, n=10) expectorated CF sputum. (Unpaired t-test, * indicates p≤0.05)

To assess the impact of Pf phages on TOB diffusion, we collected sputum samples from three pwCF who had *Pa* infections but had no Pf detected in the sputum sample by qPCR (*Pa*+Pf-). Pf status was determined using a protocol previously reported by our group (*20*) (**Table S1**). Half of these samples were spiked with Pf phage (*Pa*+Pf+). We observed that the fluorescence of TOB in the bleached region after 3 minutes was substantially lower in *Pa*+Pf+ sputum compared to *Pa*+Pf− sputum (**Figs. 2B and S11**) and the calculated apparent diffusion coefficient was also found to be lower in the case of *Pa*+Pf+ sputum (**Fig. 2C**), indicating slower TOB diffusion in the presence of Pf.

**Fig. 2.**
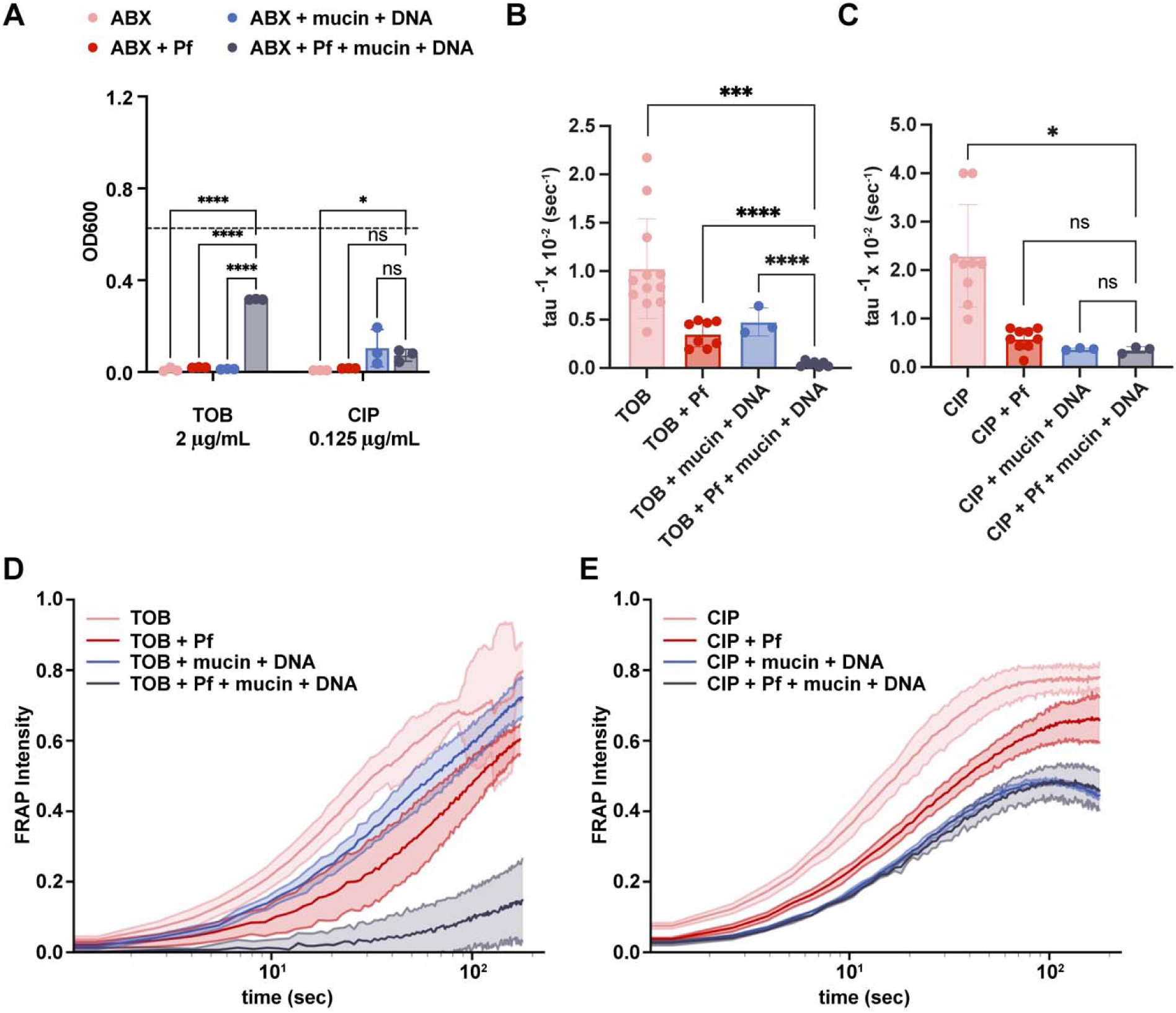
Pf and sputum polymers affect the efficacy and diffusion of tobramycin but not ciprofloxacin. **(A)** To assess the impact of filamentous bacteriophages (Pf) and lung polymers on antibiotic efficacy, we measured TOB and CIP killing of *Pseudomonas aeruginosa* (*Pa*) (n=3). antibiotics = ABX; tobramycin = TOB; ciprofloxacin = CIP. Dashed line: optical density of bacteria grown in control media. FRAP recovery curves of antibiotic diffusion were determined, and the diffusion rate, tau^−1^, was plotted for **(B, D)** Cy5-TOB in PBS (n = 12), Pf (n = 10), mucin and DNA (n = 3), and Pf with mucin and DNA (n = 7). FRAP recovery curves are also shown for **(C, E)** Cy5-ciprofloxacin in PBS (n = 9), Pf (n = 9), mucin and DNA (n = 3), and Pf with mucin and DNA (n = 3). (* indicates p≤0.05, *** indicates p≤0.001, and **** indicates p≤0.0001. (a): 2way-ANOVA for OD_600_; (c, e): Unpaired t-test for tau^−1^ plots.)

### Pf phages inhibit the diffusion and efficacy of TOB in sputum polymers

We then sought to interrogate how Pf impacts the diffusion and efficacy of TOB in more controlled systems. To this end, we studied a set of antibiotics against *Pseudomonas aeruginosa* PA14: TOB, colistin (COL), ciprofloxacin (CIP), and aztreonam (AZT). Because sputum is highly heterogenous and variable from patient to patient, to facilitate reproducible mechanistic analyses of phage/polymer/antibiotic interactions, we made artificial sputum by combining 4 mg/mL DNA and mucin to 8% solids (w/w) for these studies (*27*). Notably, we did not include the amino acids, lipids, egg yolk, etc. that are often incorporated into artificial sputum preparations (*37*) to primarily focus on the interaction between sputum polymers and Pf.

We first examined the impact of Pf phages and sputum polymers on bacterial killing. We found that the combination of Pf phages and sputum polymers resulted in a significantly decreased bacterial killing for TOB (**Fig. 2A**) and COL (**Fig. S8A**) but not for CIP (**Fig. 2A**) and AZT (**Fig. S8A**). We then used FRAP to quantify the diffusion rate of cy5-TOB and cy5-CIP in sputum polymers. In **Fig. 2D** and **Fig. 2E**, the fluorescent recovery curve is shown for cy5-TOB and cy5-CIP, respectively. The effective diffusion rate, found by assuming the fluorescent recovery is due purely to Fickian diffusion, is given by tau^−1^ (with units sec^−1^), whose value for each condition is plotted for cy5-TOB and cy5-CIP in **Fig. 2B** and **Fig. 2C**, respectively. Comparing the effective diffusion rates of TOB and CIP with the condition of having both Pf phages and artificial sputum present, we found that the combination of Pf and artificial sputum resulted in a significantly decreased fluorescence recovery for TOB (**Fig. 2B, D**) but less so for CIP (**Fig. 2C, E**). Additional comparison statistics support that TOB exhibited sharper drops in diffusion rate due to Pf and artificial sputum (**Fig. S10A, B**). Together, these data suggest that Pf and sputum polymers (DNA and mucin) have distinct effects on the diffusion of different antibiotics.

### Isothermal titration calorimetry results support electrostatics as the driving force in TOB sequestration by Pf phages and sputum biopolymers

To interrogate the mechanism underlying decreased antibiotic efficacy in the presence of Pf phages, we assessed the interaction strength between Pf and antibiotics using isothermal titration calorimetry (ITC). Because TOB is a highly positively charged molecule at physiological pH (**Table S2**) (*38*), we hypothesized that negatively charged Pf and sputum polymers would reduce the diffusion of positively charged antibiotics due to electrostatic attraction, diminishing antimicrobial efficacy. For these assays, we used only Pf and DNA to focus on this specific interaction in the measurement. TOB or CIP was titrated into solutions of either Pf phages, DNA, or both. The heat released from titrating in TOB to each solution was mostly positive, which corresponds to an energetically favorable interaction between TOB and Pf and DNA (**Fig. 3A**). On the other hand, the heat released from titrating CIP into each solution was mostly negative (**Fig. 3B**), indicating energy was consumed and the mixing is energetically unfavorable.

**Fig. 3.**
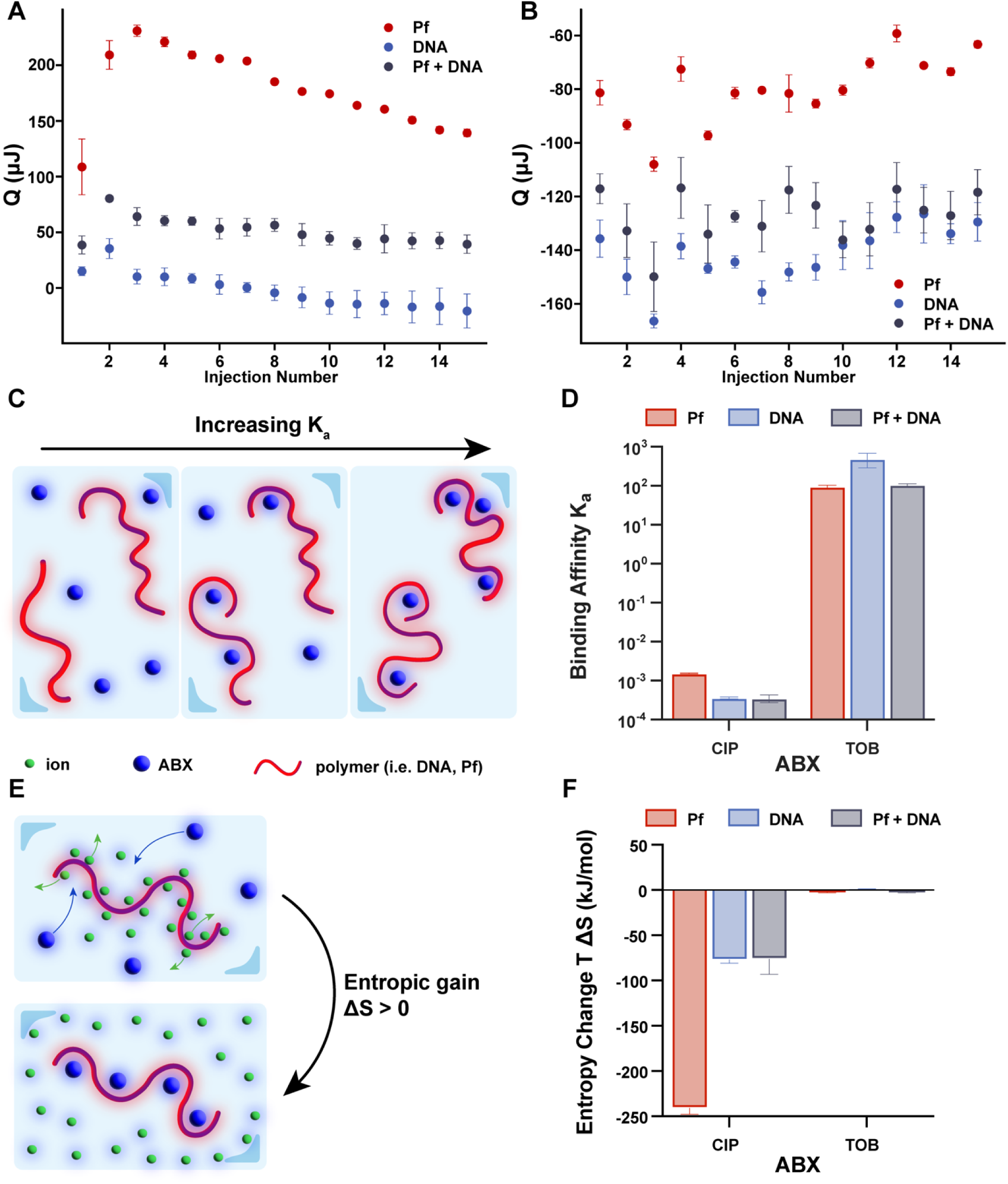
Isothermal titration calorimetry reveals that the electrostatic interaction is antibiotic specific. **(A)** Heat per injection corrected for blank conditions for three conditions: tobramycin (TOB) in 4 mg/mL DNA (n = 3), TOB in 10e11 pfu/mL Pf4 (n = 3), and TOB in 4 mg/mL DNA and 10e11 pfu/mL Pf4 (n = 3). **(B)** Heat released per injection corrected for blank conditions for three conditions: ciprofloxacin (CIP) in 4 mg/mL DNA (n = 3), CIP in 10e11 pfu/mL Pf4 (n = 3), and CIP in 4 mg/mL DNA and 10e11 pfu/mL Pf4 (n = 3). **(C)** A larger binding affinity, K_a_, indicates more antibiotics and polymers bound together at equilibrium. **(D)** Binding affinities for the positively-charged antibiotic, TOB, and neutrally charged antibiotic, CIP, in Pf phage (TOB: n = 2; CIP: n = 2), DNA (TOB: n = 3; CIP: n = 2), and both Pf phage and DNA (TOB: n = 2; CIP: n = 3) are shown. Error bars show +/− 1 standard deviation. **(E)** Larger and more positively-charged antibiotics displacing positive counter ions surrounding negatively-charged polymers like DNA and Pf phage can lead to a positive change in entropy. **(F)** Entropic change is positive for the positively-charged antibiotic, TOB, and negative for the neutral antibiotic, CIP. The reference concentration to determine entropy change was 0.003 mol/L.

To quantitatively derive the strength of interaction between antibiotics and polymers from the isotherms, we fit a thermodynamic model to the isotherms that accounts for thermodynamic contributions from excess entropy associated with electrostatic interaction and enthalpy of phase changes such as coacervation. The model is based on basic polymer physics principles and is described in more detail in the Methods section. This model has previously been used to analyze binding mechanisms in polyelectrolyte solutions composed of equal concentrations of oppositely charged polymers, which will first pair up electrostatically in solution before phase separating into coacervates (*39*). In our case, the assumption is that electrostatic attraction between positively charged antibiotics and negatively charged Pf and DNA would drive the process of ion pairing. If enough charged species bind together past a certain threshold, there could be phase separation of the complex of antibiotics, Pf, and DNA out of the solution. Indeed, cloudiness was observed in the chamber after titrating tobramycin into Pf and DNA, indicating a phase change had occurred (**Figure S9**).

Using the model, the binding affinity and the enthalpy change of the ion pairing process were determined. The binding affinity, K_a_, gives insight into the strength of interaction between the antibiotic and polymers in the solution, with higher values indicating that more antibiotics and polymers are bound at equilibrium (**Fig. 3C**). Consistent with our predictions, the positively charged TOB exhibited much greater binding affinity with DNA, Pf, and the DNA+Pf mixture than the neutrally charged CIP in each condition (**Fig. 3D**).

Additionally, the binding affinity can be used to calculate free energy change due to the ion pairing process, composed of energetic and entropic contributions. The energetic contribution is a fitted parameter, ΔH_IP_. It can be subtracted from the free energy change to reveal the entropic contribution to the ion pairing process, where a more positive entropic contribution correlates to more binding (**Fig. 3E**). We observed a greater negative entropic contribution to the free energy change for neutrally charged CIP than the positively charged TOB (**Fig. 3F**). The more positive entropic change due to the ion pairing between oppositely charged TOB and sputum components (Pf phages, DNA) compared to CIP is in line with what has previously been found in oppositely charged polyelectrolyte mixtures (*39–41*), suggesting that TOB interacts with sputum components in a charge-dependent manner. Entropy change values for DNA and several multivalent ions Cu^2+^, Co^2+^, Ni^2+^ were found to lie within the range of −12.8 cal/molK to 10.8 cal/molK (*42*). For TOB binding to DNA, we found that the entropy change is 12.56 cal/molK, which is just above but comparable to the range that previous work found for DNA and the above three multivalent ions. The higher entropy value indicates that TOB binding to DNA is even more favorable than the above mentioned ions and is a largely entropically driven process.

Together, the data indicates that TOB but not CIP binds to both Pf phages and DNA and implicates electrostatic interactions as the driving force. Notably, we did not observe substantial differences in the binding affinities between TOB in Pf alone and in the Pf+DNA mixture. This suggests that any higher order interactions between Pf phage and DNA (e.g. the ability of Pf to organize DNA into a crystalline network) do not impact binding strength.

### DNA and Pf phages hinder the diffusion of TOB in a pH-dependent manner

We sought to assess the impact of pH in these interactions. Because pH changes could be expected to alter a molecule’s effective charge, altering pH allowed us to assess the impact of electrostatic forces. We repeated the bacterial killing assay in Figure 2, using only DNA and Pf phage. We found that mixtures of TOB and DNA but not CIP and DNA reduce bacterial killing (**Fig. 4A**). We then repeated these studies over a range of pH values. Bacterial growth was not affected until the pH was above 10. We found that the efficacy of TOB was restored as the pH increased above 7 (**Fig. 4B**) because the amide groups in tobramycin at higher pH values undergo deprotonation (*43*), thereby decreasing the positive charge on the molecule. The pH dependence of TOB antimicrobial activity indicated that the charged interactions between Pf phage, DNA, and TOB impact TOB efficacy. In contrast, the antimicrobial activity of CIP was not significantly affected by pH (**Fig. 4C**).

**Fig. 4.**
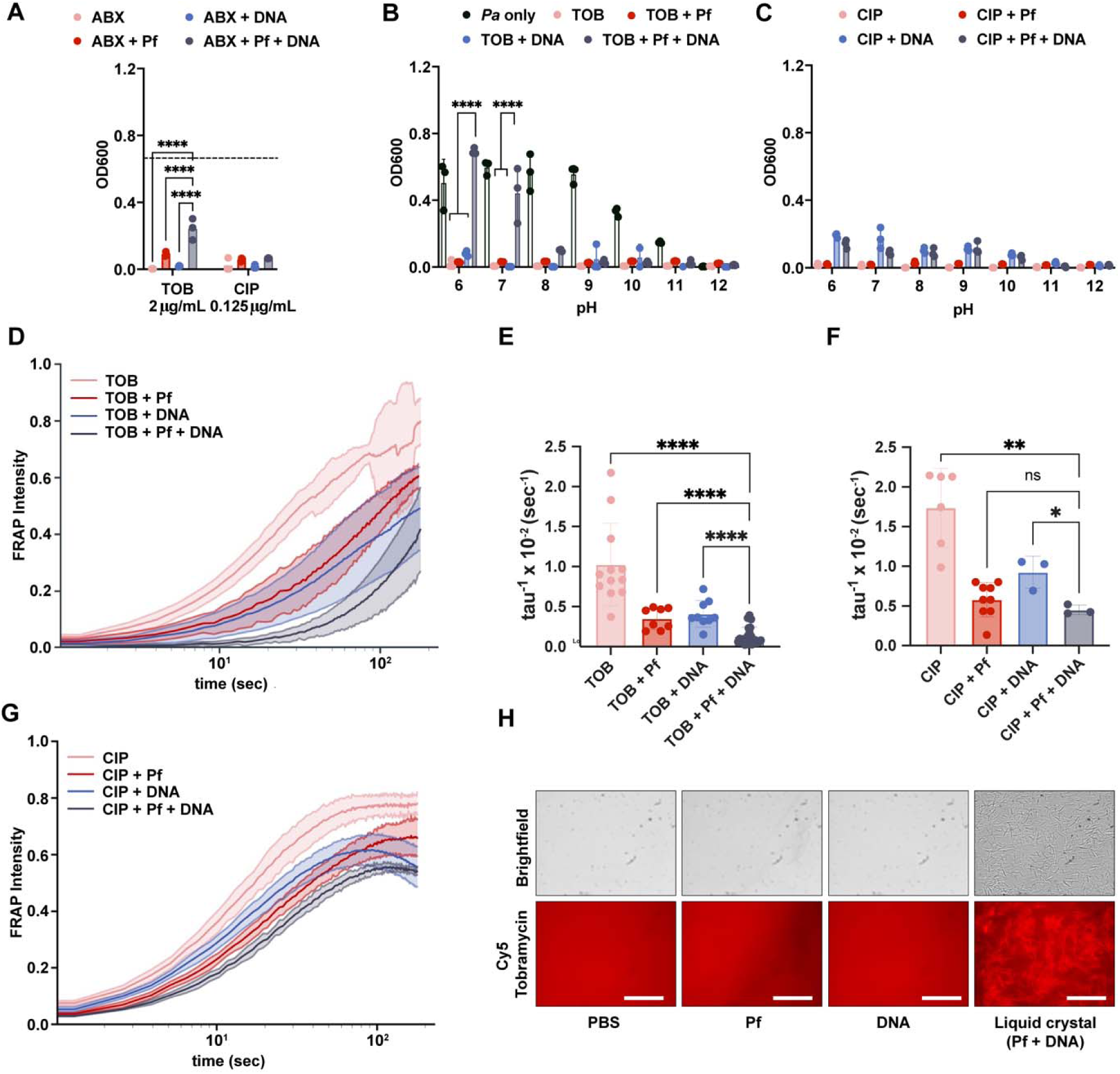
Sputum polymers mixed with Pf phage reduce efficacy and diffusion of antibiotics in a charge-dependent manner. **(A)** Pf and DNA affect TOB and COL efficacy against *Pa.* Dash line indicates the OD for *Pa* alone (n=3). antibiotics = ABX; tobramycin = TOB; colistin = COL; ciprofloxacin = CIP; aztreonam = AZT. Phages and DNA affect **(B)** TOB, but not **(C)** CIP, antimicrobial activity in a pH-dependent manner against PA14 (n=3). **(D)** The fluorescence intensity from FRAP experiments of Cy5-TOB in PBS (n = 12), Pf (n = 9), DNA (n = 12), and both Pf and DNA (n = 13) are shown. The effective diffusion rate, tau^−1^, was found from the FRAP recovery curves and plotted for **(E)** Cy5-TOB and **(F)** Cy5-CIP. **(G)** FRAP recovery curves for Cy5-CIP in PBS (n = 9), Pf (n = 9), DNA (n = 3), and both Pf and DNA (n = 3) are shown. **(H)** The binding of Cy5-TOB to Pf phages-mediated liquid crystals (DNA+Pf condition) was visualized by fluorescent microscopy. (* indicates p≤0.05, ** indicates p≤0.01, and **** indicates p≤0.0001. 2way-ANOVA for OD_600_; Unpaired t-test for tau^−1^ plots.).

In parallel, FRAP experiments were performed to determine the diffusion of antibiotics through DNA, Pf phages, and mixtures of DNA and Pf phages at pH 7. As expected based on previous studies, we observed colocalization of Cy5-TOB within the liquid crystal tactoids formed in mixtures of DNA and Pf phages (**Fig. 4H**) (*27*). The fluorescent recovery decreased as more components were added to the solution (**Fig. 4D**). Cy5-TOB (pink curve), by itself, recovered fully. Adding Pf phages (red curve) or DNA (blue curve) slowed the effective diffusion rate, tau^−1^, but combining both (black curve) slowed the diffusion even further (**Fig. 4E**). The diffusion of cy5-CIP was not affected by the presence of both Pf and DNA (**Fig. 4F-G**). Together, the results for bacterial killing at varying pH and FRAP curves implicate electrostatic forces in TOB affinity for and inactivation by Pf phage and DNA.

We also interrogated the alternative hypothesis that increased quantities of macromolecules were decreasing the mesh size to be smaller than the size of the antibiotic molecule and limiting diffusion by being a physical barrier. To test this, we used FITC-dextran molecules of the same order of magnitude in size as TOB to investigate whether changes in mesh size would limit antibiotic diffusion. We found that the mesh size did not change due to the addition of Pf and DNA, as the FITC-dextran molecule exhibited similar diffusion rates in all conditions with sputum components (**Fig. S3**).

### In modeling studies, higher-order structures formed by DNA and Pf phages reduce the diffusion of charged antibiotics through electrostatic interactions

We sought to understand better the binding interactions that underlie TOB sequestration through modeling studies. To model this transient binding that can occur and its effect on fluorescent recovery in a FRAP experiment, we derived a 2-state model that assumes that the antibiotic molecule transiently switches between a free state and being bound to another constituent (**Fig. 5D**). The model contains binding and unbinding rates, which govern the kinetics of the associations, as well as a parameter α, which is the ratio of the diffusion in the bound state to the diffusion in the free state. A detailed derivation and description of this model can be found in the Methods section. The 2-state model was fit to FRAP data for both TOB (**Fig. 5A**) and CIP (**Fig. 5B**) diffusion in Pf phages to determine the kinetics of the binding to Pf phages, and the relative decrease in diffusion related to binding (**Fig. 5A-B**, blue curves). We similarly fitted the 2-state model to our FRAP data for TOB and CIP diffusion in DNA alone (**Fig. 5A-B**, red curves).

**Fig. 5.**
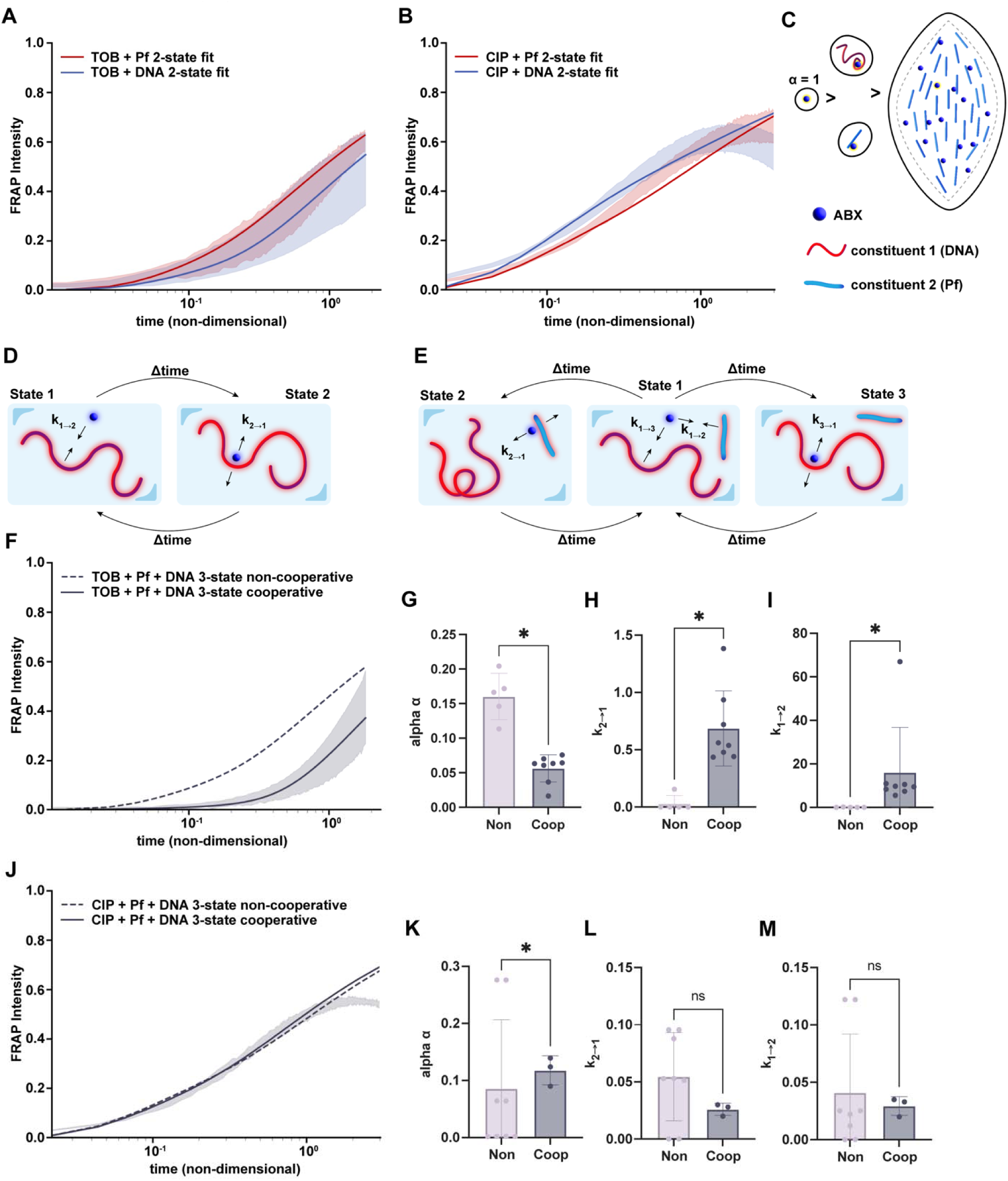
A set of 2-state and 3-state models demonstrates that liquid crystals reduce the diffusion of charged antibiotics through electrostatic interaction. The distribution over one standard deviation of the FRAP recovery data is shown along with the best fit of the 2-state model to each condition for **(A)** Cy5-TOB and **(C)** Cy5-CIP in either Pf phages or DNA. **(C)** Schematics demonstrate that electrostatic binding could affect the diffusion of positively-charged antibiotics, such as TOB, by increasing the size of the diffusing mass. **(D)** The FRAP data were modeled using a newly derived diffusion model that accounts for electrostatic attraction between the antibiotic and another constituent (2-state model). **(E)** The FRAP data were modeled using a newly derived diffusion model that accounts for electrostatic attraction between the antibiotic and 2 other constituents, attributing different diffusion times to each type of bound state: antibiotic bound to Pf phages, antibiotic unbound to anything, and antibiotic bound to DNA. The distribution over one standard deviation of the FRAP recovery data is shown along with the non-cooperative and cooperative 3-state models to each condition for **(F)** Cy5-TOB and **(j)** Cy5-CIP in mixtures of Pf phages and DNA. The non-cooperative FRAP recovery assumes no additional contribution to diffusion due to liquid crystal (dashed), while the cooperative FRAP recovery shows the effect of liquid crystal formation on antibiotic diffusion (solid). The change in the diffusion and binding parameters found via the model for **(G-I)** TOB or **(K-M)** CIP bound to Pf phages or antibiotic bound to a liquid crystal structure of Pf phages and DNA are shown. **(G-I)**The values for TOB show statistically different values demonstrating an additional diffusion hindrance from the liquid crystal. Alpha is the change in relative diffusion compared to antibiotic alone in a buffer solution, where a lower alpha value corresponds to a slower diffusion. k_2→1_ and k_1→2_ are unbinding and binding rate constants, respectively, where the ratio of the two gives the equilibrium constant of the binding interaction. A higher equilibrium constant implies a longer time spent bound. (* indicates p≤0.05)

**Fig. 6.**
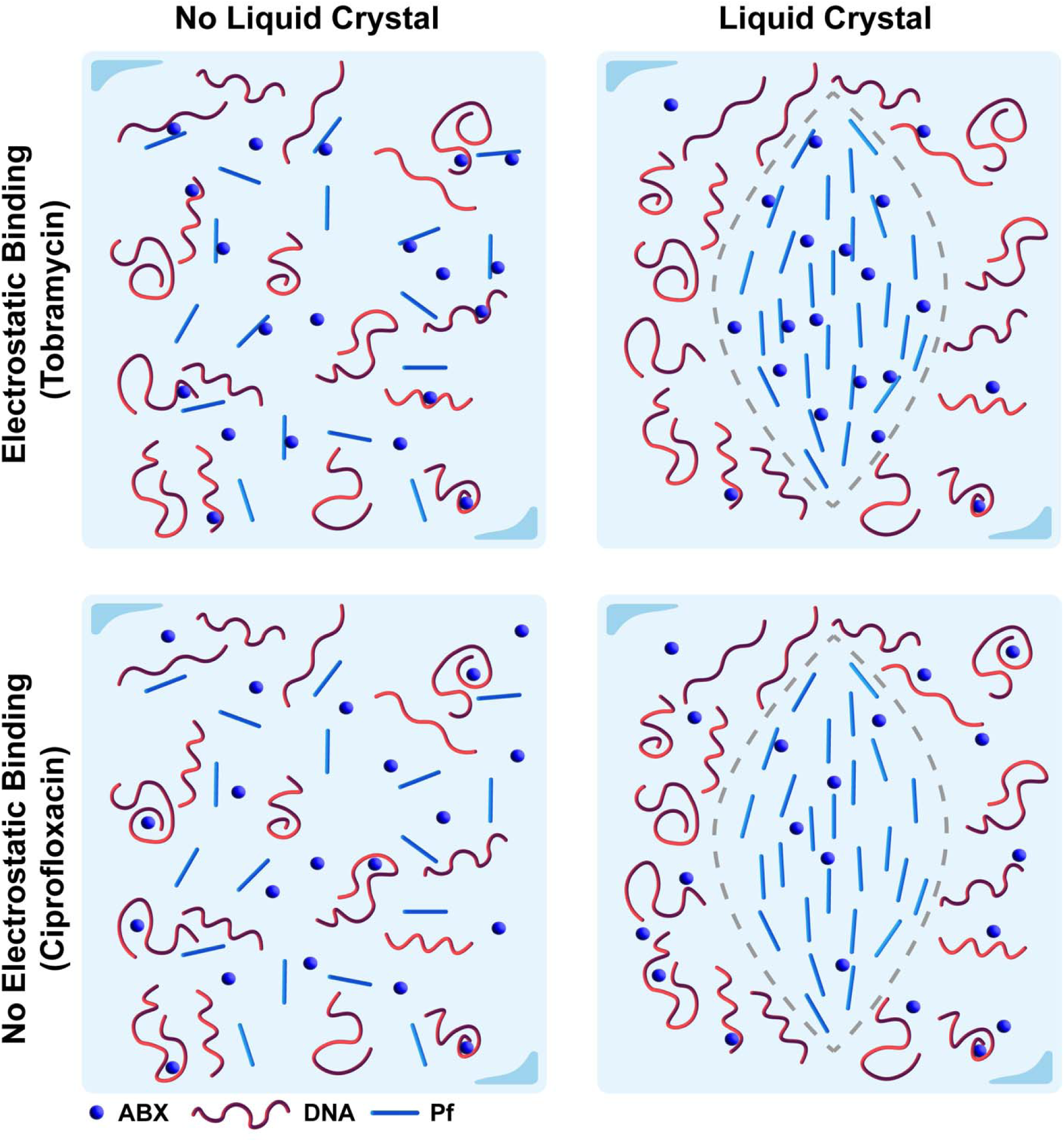
Depiction of the interaction mechanism among positively-charged and neutral antibiotics with the sputum components Pf phages and DNA. For positively-charged antibiotics like TOB, the lack of a liquid crystal formation hinders diffusion through binding with sputum components Pf phages and DNA (top left). When there is liquid crystal formation, these antibiotics are further sequestered and experience increased rate of binding due to the higher density of polymers in the liquid crystal tactoid (top right). For neutral antibiotics like CIP, the lack of electrostatic attraction results in little influence of sputum polymers on diffusion, though in the liquid crystal the higher density of polymers will decrease effective diffusion (bottom row).

In the case where both Pf phages and DNA are present in the solution, we hypothesized that the diffusion of antibiotics should be a function of antibiotics in three states: free diffusion, bound to Pf phages, and bound to DNA (**Fig. 5E**). To model this scenario, we derived the 3-state model that includes kinetic parameters to govern the transitions among these three states and ratios of the relative diffusion rates in each state (described in detail in the Methods). In this non-cooperative 3-state model, we assume that the binding of antibiotics to DNA is independent of the binding of antibiotics to Pf phage. Thus, we can use the parameters we derived for antibiotic diffusion in DNA alone and Pf phages alone from fitting the 2-state model to these scenarios to generate a predicted diffusion curve with this non-cooperative 3-state model. We found that the resulting non-cooperative 3-state model predicted a faster recovery in TOB fluorescence than the experimental data showed (**Fig. 5F**). On the other hand, the fluorescent recovery of CIP in DNA and Pf phages was almost perfectly predicted by the non-cooperative 3-state model (**Fig. 5J**). Given previous findings of liquid crystal formation between Pf phages and DNA, we decided to quantify the effect of liquid crystals on diffusion in these conditions. To do so, we used the same 3-state model, but this time, we only used the parameters for DNA obtained from the 2-state model fit of the FRAP data for antibiotics in DNA. The parameters for the second constituent to which the antibiotics can bind were fit to the FRAP data for a mixture of DNA and Pf phages. By fitting the parameters of the second constituent to the FRAP data, we imply that there is cooperativity between antibiotic binding to DNA and binding to Pf. Thus, these new parameters for the second constituent reflected any additional hindrance to antibiotic diffusion from DNA and Pf phage interactions, which stem from liquid crystal formation. For TOB, the diffusion rate α of the second constituent was lower than the α in the non-cooperative 3-state model (**Fig. 5G**). In contrast, the diffusion rate for CIP remained, on average, the same between the non-cooperative and cooperative models (**Fig. 5K**). The kinetic parameters k_1-2_ and k_2-1_ for TOB suggested faster binding to the liquid crystal and more time spent bound to the liquid crystal than when TOB was in a solution of just Pf phages (**Fig. 5H-I**). The kinetic parameters for CIP were similar between the cases of Pf phages only (non-cooperative) and Pf+DNA liquid crystals (cooperative) (**Fig. 5L-M**). Overall, we found that for TOB, not only was the diffusion in Pf phage and DNA slower than when we assumed non-cooperativity, but cooperativity also appeared to increase the rate of binding of TOB to the second constituent (**Fig. 5 G-I**).

To further validate the effect of the liquid crystal on TOB diffusion, we compared the FRAP data for TOB in DNA and Pf phages to data for TOB with DNA and a non-liquid crystal forming bacteriophage, LPS5 (**Fig. S4**). Combining DNA and LPS5 did not significantly change the FRAP recovery compared to TOB diffusion in DNA alone (**Fig. S4A-B**). Using the same 3-state FRAP models above, we found that the predicted FRAP recovery with the 3-state non-cooperative model aligned with the FRAP recovery curve for TOB in a mixture of DNA and LPS5 (**Fig. S4A-B**). Furthermore, we fitted the 3-state non-cooperative model to the FRAP recovery data for the condition of DNA and LPS5. We found no significant difference in the parameters between the cooperative and non-cooperative models (**Fig. S4C**). The main difference between LPS5, a lytic phage that infects *Pa*, and Pf phage is the formation of liquid crystals between Pf and DNA, so the results suggest that liquid crystals significantly impact TOB transport.

## Discussion

In this study, we have asked how Pf phages and polymers present in sputum prevent the diffusion of TOB, an inhaled antibiotic commonly used to treat chronic *Pa* infection in pwCF. We demonstrated that the diffusion of TOB, a positively charged antibiotic, through networks of Pf and sputum polymers, is limited by electrostatic interactions. Conversely, CIP, a neutrally charged antibiotic, exhibits limited electrostatic attraction with Pf or sputum components. Moreover, higher-order, crystalline structures formed between Pf phages and sputum polymers further slow the diffusion of TOB due to these charge-based interactions. These results are consistent with previously published studies showing that positively charged antibiotics have reduced antimicrobial activity when Pf is present (*27, 34*) and establish a biophysical explanation for these interactions.

The data presented here generally agree with previous modeling studies. Rossem *et al.* modeled antibiotic diffusion in liquid crystal tactoids formed by Pf phages and sputum biopolymers (*36*). Their mathematic model yields results for antibiotic diffusion that agree broadly with the experimentally observed results. However, their model constrained the problem to the tactoid volume, whereas in real sputum environments, most of the volume is unlikely to consist of liquid crystals. The focus on the tactoid space could explain Rossem *et al.*’s own observation that the timescales predicted by their model are not sufficiently long to explain antibiotic tolerance on the order of hours (*27*). Likely in the sputum environment, liquid crystal tactoids form spontaneously due to local density changes in biopolymers and Pf phages so that the sputum environment contains regions of tactoids and regions devoid of tactoids. Our focus when modeling antibiotic diffusion was on handling the temporal heterogeneity observed through the binding and unbinding of the sputum environment and not just the tactoid volume. The larger scope of our model sacrifices spatial resolution, which is captured explicitly in Rossem *et al.*’s approach. Thus, a comprehensive model would incorporate aspects of both our model and that of Rossem *et al*.

Our own models reveal cooperativity between the binding of TOB to Pf phages and DNA but not between the binding of CIP to Pf phages and DNA. Despite the fact that the pH measurement and how pH affects antimicrobial in CF airways have been inconclusive, several studies have suggested that the average pH in sputum and airway is 6.6-7 (*44, 45*), which is the condition that tobramycin carries positive charges. If the binding between TOB and sputum components were the only factors hindering its diffusion, we would not have observed this cooperativity. Thus, cooperativity arises from the additional influence of liquid crystals that form at the DNA and Pf phage concentrations used in our experiments (*27*). The set of models developed in this paper provides powerful tools for analyzing diffusion with binding due to how the parameters can be translated across the three different models. The simplicity behind these models makes them easily applicable to a wide range of scenarios where one would want to capture the main contributing factor influencing observed diffusion trends.

Further modeling studies need to be performed to better understand if the structure of the polymer or charge of the polymer plays a larger role in hindering antibiotic diffusion in liquid crystals and whether the results we observed for DNA can be extended to other negatively charged biopolymers found in the sputum and *Pa* biofilms. For example, mucin is a large component of the secretory apparatus in the airways, so any interactions between mucin and Pf would be valuable for therapeutic purposes. In this study, we focused on DNA due to its drastically increased presence during *Pa* infection compared to other polymers like mucin. Our data shows that Pf with artificial sputum (containing mucin) correlates to reduced antibiotic diffusion (**Fig 2**); however, it necessitates the study of Pf and mucin interaction as a next step. Similarly, we confined our study to TOB and CIP. Aerosolized AZT is also available for chronic *Pa* infection in pwCF in the United States. It would be therefore important to extend these studies to other antibiotics in the future.

Electrostatic binding between Pf phage and positively charged antibiotics could be attenuated as a strategy for addressing antibiotic diffusion hindrance. For example, positively charged antimicrobial peptides, such as LL-37, have been shown to interact with negatively charged cellular components (*46*), such as lipopolysaccharides (*47*), eDNA (*48*), and dsRNA (*49*). This molecule and other similarly charged molecules could compete with positively charged antibiotics in binding to Pf phage and sputum biopolymers. By replacing the antibiotics in the bound state, more antibiotics can freely diffuse, thus potentially increasing antibiotic efficacy.

One limitation of this work is the simplified model system we used because it lacks some of the complexities of sputum produced in the CF airway. We nonetheless believe that our models capture some of the relevant biophysical attributes of this system. It has previously been shown that high molecular weight DNA with Pf phages alone can lead to the formation of liquid crystal tactoids (*27*). Studies have suggested that DNA is an important structural component in biofilms and is negatively associated with pulmonary function and disease severity in pwCF (*50–52*). Therefore, we simplified our model system to just Pf phage and DNA. Future studies will examine antibiotic and Pf phage interactions in more complex sputum formulations.

In summary, our analysis suggests that Pf phages in sputum reduces the diffusion of charged antibiotics due to a greater binding constant. In the future, we will leverage this understanding of the interaction between positively charged antibiotics and Pf phages to design or discover molecules that recover the diffusion of and rescue the antimicrobial activities of positively charged antibiotics against the clinically significant *Pa*.

## Material and Methods

### Collection of CF sputum

Sputum samples from patients with CF were collected by spontaneous expectoration during clinic visits or hospitalizations. Patients and healthy controls were consented for sputum collection under IRB approved protocols #37232, 56996, or 11197. Sputum was aliquoted and frozen at −80 L prior to use.

### Bacterial culture

*Pseudomonas aeruginosa* PA14 was streaked from a frozen glycerol stock onto LB agar (, BD BBL) plates and incubated overnight at 37 L with 5% carbon dioxide (CO_2_) until individual colonies formed. A single colony was inoculated into 3 mL LB media and grown at 37 L in a shaking incubator at 250 rpm to an OD_650_ of 0.4, corresponding to ~5×10^8^ CFU/mL. Bacteria cultures were adjusted to 1×10^6^ CFU/mL to prepare a working stock for all experiments.

### Bacteriophage isolation and propagation

Phages were propagated based on the previously described protocol (*53*). Briefly, planktonic bacterial cultures were grown to an OD_650_ of 0.4, then infected with phage stock and further grown for 2 hours. Then, 150 mL of LB media were inoculated with phage-infected bacterial cultures and incubated at 37 °C overnight shaking. The overnight cultures were centrifuged (8000 × g, 30 min, 4 °C), and filtered through a 0.22 μm PES membrane (Corning, Corning, New York, Product #4311188) to remove bacterial debris. The filtrate was treated with 5 U/mL benzonase nuclease (Sigma-Aldrich, Saint Louis, MO, Catalog #E8263) at 37 °C. After benzonase treatment, filtrates were centrifuged (8000 × g, 4 °C) overnight, and the pellet was resuspended and quantified for titer by plaque assay.

### *In vitro* antimicrobial activity

Four antibiotics were used in this experiment, including TOB, CIP, COL, and AZT. Each antibiotic was dissolved in sterile distilled water to make stock solutions with concentrations of 1 mg/mL, and diluted to indicated concentration in LB. Salmon sperm DNA (Sigma-Aldrich, Saint Louis, MO, Product #D1626) and Mucin (Sigma-Aldrich, Saint Louis, MO, Product #M1778) were dissolved in LB. The bacterial killing by different antibiotics in various conditions was determined according to the standard Clinical and Laboratory Standards Institute (CLSI) broth-microdilution method and adapted from previous studies (37). Briefly, a 100 μL aliquot of a bacterial suspension was added to each well (n=3) containing 100 μL solutions that contained antibiotics only, antibiotics with Pf phages, antibiotics with DNA or DNA added mucin, or antibiotics with Pf with DNA or DNA added mucin in a 96 well plate to achieve 5×10^5^ CFU/well. The final concentrations of Pf phages, DNA, and Mucin are 10^11^ pfu/mL, 4 mg/mL, and 8% solids, respectively. The 96-well plate was incubated at 37 for 18 - 24 hours under static conditions. The OD600 was measured in a Biotek Synergy HT (BioTek, Winooski, VT, USA), where higher OD600 indicated less bacterial death.

### Isothermal titration calorimetry (ITC) studies

The thermal behavior of antibiotics interaction with Pf phages and polymers was studied using the ITC instrument (TA Instruments Nano ITC). Antibiotics were titrated from the syringe into the sample cell that contains PBS only, 10^11^ pfu/mL Pf4 phages in PBS, or 10^11^ pfu/mL Pf4 and 1 mg/mL DNA in PBS. After each injection, the heat exchange produced from the interaction was measured by ITC calorimeter. Between each injection, the system was allowed to attain equilibrium.

For an experiment titrating component 1 (i.e. antibiotic) into component 2 (i.e. DNA), data was obtained from for the 1) titration of component 1 into component 2, 2) titration of component 1 into a buffer solution without component 2, and 3) titration of a buffer solution without component 1 into component 2. The last two titrations are considered blanks for the first titration and are subtracted from the first titration to correct for entropic change due to volume change. This corrected heat change data was fit using a non-linear least squares regression to a two-component thermodynamic binding model. The binding model contains two contributions: one from ion pairing (“IP”), or electrostatic binding, and one from phase change (“C”), possibly precipitation or liquid crystallization (*39*).

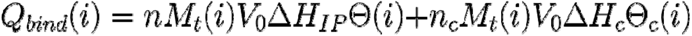

For antibiotic binding to a polymer, there are multiple binding sites per polymer chain. Each binding event of an antibiotic molecule to the polymer is assumed to be independent of other binding events along the same polymer.

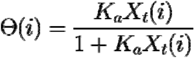

The fitted binding affinity, K_a_, was directly compared across the conditions. The entropy change, ΔS, was found by the following thermodynamic relation using a reference concentration of 0.003 mol/L, which is about the concentration of CIP used in its ITC measurements. In this equation, ΔH_IP_ and K_a_ are found through fitting the data to the model, and R and T, the gas constant and the temperature, are known quantities.

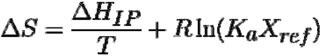

### Measurements of diffusion of fluorescently tagged antibiotics by fluorescent recovery after photobleaching (FRAP)

Cy5 labeled antibiotics (TOB and CIP) purchased from Biosynthesis Inc. were used in the experiment. Six conditions were used to investigate the diffusion of antibiotics, including control, Pf4 phages only, polymers only, and Pf4 mixed with polymers. Each condition was added with 1 μL of fluorescent antibiotic with concentration of 1 mg/mL. FRAP experiments were performed using a Leica DMI 8 confocal microscope with a 10x dry objective. In each FRAP experiment, a pre-bleach image was acquired at 2% of the maximum laser intensity. Then, the mixture was bleached for 30 seconds at the maximum laser intensity and recorded intensity recovery for 3 minutes.

### Methods for FRAP Analysis

A Green’s function was used to denote the distribution of trajectories of the fluorescent molecules in a 1-D system and combined with systems of differential equations accounting for the kinetics of binding events to model the fluorescent recovery when binding impacts diffusion. We assume no photofading. The three models presented here, 1-state, 2-state, and 3-state models, allow hypothesis testing of any type of binding interactions that could be occurring in a FRAP experiment and allow extraction of more information from a single FRAP measurement.

### 1-state FRAP model: Diffusion only

1-state model assumes that fluorescent particles move with diffusion coefficient D with no kinetic events or binding occurring. A 1-D line of length L is imagined, with the position x = 0 at the center such that from x = 0 to the edge of the line is a distance L/2 (**Figure S5**). It is assumed that the probability that a particle starting at a position x_0_ ends up at a position x at time t follows a normal distribution.

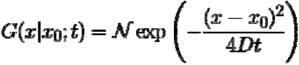

In this system, numerous fluorescent particles are in the area, but the fluorescence at position x = 0 is measured as the signal. The signal at position x = 0 is found by integrating over all possible starting particles at positions x_0_ = 0.1L, 0.2L, 0.3L… that will be at position x = 0 at time t.

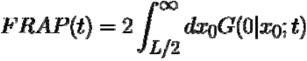

After replacing the probability distribution in the above equation with the normal distribution function, the fluorescent intensity as a function of time for an area of interest with size L is found.

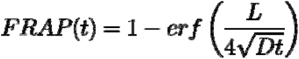

Furthermore, this model can be nondimensionalized with nondimensionalization factor D/L^2^.

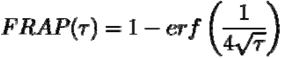

### 2-state FRAP model: Diffusion and binding to one constituent

The second model, 2-state model, assumes that in addition to the diffusion of the fluorescent particle at some diffusion coefficient D_1_ = D (“State 1”), these particles can bind to some other constituent that has a diffusion coefficient of D_2_ = αD (“State 2”), where α < 1 (i.e. being bound to this other constituent slows down the particle’s diffusion compared to when it is free). The binding process is transient, with a rate of binding to the other constituent k_1 2_ and unbinding from the other constituent k_2 1_ (**Figure 5D**). The following reaction and defined kinetic parameters describe this 2-state scenario.

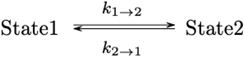

It should be noted that from here on the use of the variable t corresponds to nondimensionalized time, which is denoted in the 1-state model derivation with the letter τ.

A Green’s function is used to denote the spatial integration of the fluorescence as in the 1-state model. It is assumed an initial condition for the fluorescent signal:

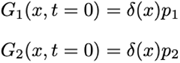

where

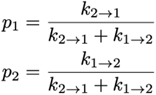

With two states, this model requires a set of differential equations to describe the diffusion and kinetics of the system.

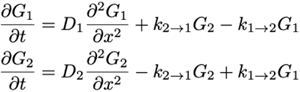

Solving this set of equations involves Laplace (t → s) and Fourier (x → k) transforms to make the operation algebraic. The Green’s functions for state A and state B, which represent the fluorescent contributions from each state to the overall average fluorescent intensity, are solved for and summed to get the total of the fluorescent signal because the total fluorescence comes from free particles in solution and particles bound to polymers. Both states contribute to the total fluorescence but will simply have different rates of fluorescent fluctuation given the differences in diffusion. Lastly, the transform is inverted to give the fluorescent signal as a function of time. The 2-state model was evaluated using diffusion rates for the two states that are equal and found to agree with the 1-state model using that same diffusion rate, which is as expected (**Figure S6A**). When State 2 is given a slower diffusion rate and the rate of unbinding is lowered, the 2-state model predicts a slower recovery before eventually returning to the same full fluorescent recovery as the 1-state model (**Figure S6B**).

### 3-state FRAP model: Diffusion and binding to two kinetically distinct entities

In this scenario, there are two different entities that the particle can bind to, and those two entities also have different, distinct diffusion rates (**Fig. 5E**). “State 1” is still the free particle state. “State 2” is when the particle is bound to the first constituent and has diffusion D_2_ = α_2_D. “State 3” is when the particle is bound to the second constituent and has diffusion D_3_ = α_3_D. In both cases the diffusion of the bound state is lower than that of the free particle, denoted as D_1_ = D (i.e. α_2_ and α_3_ < 1).

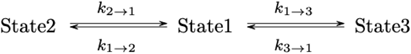

The binding and unbinding of the particle to each constituent is governed the respective on and off rates. Similarly to the 2-state model, a set of differential equations governs the diffusion and kinetics of this system with initial conditions for the Green’s functions.

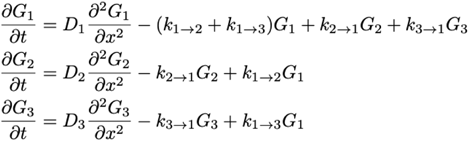

Initial conditions are similar to the 2-state model.

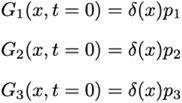

where

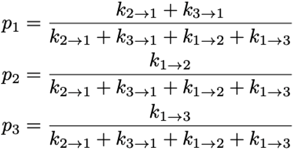

Solving the set of differential equations involves Laplace and Fourier transformations to obtain the individual Green’s functions for each state. The inversion of the Laplace transform to the time domain gives the fluorescent recovery over time.

The 3-state model was evaluated using diffusion rates for the two bound states that are equal to the free state and found to agree with the 1-state model and 2-state model in the limit of equal diffusion rates, which is as expected (**Figure S7A**). When we slow the diffusion of States 1 and 2, the 3-state model predicts a slower recovery and exhibits a delay in the recovery from even the 2-state model, which has the same diffusion rate as one of the states in the 3-state model. Thus, the 3-state model can account for the additional slowdown of the fluorescent particle when it is bound to an even slower additional constituent (**Figure S7B**).

### Fitting the FRAP models to the FRAP data

FRAP recovery curves were fit to the FRAP models using nonlinear least-squares regression. FRAP data for antibiotics diffusing in buffer, where there is assumed to be no binding occurring, was fit to the 1-state model. The nondimensionalization factor from this fit was used to nondimensionalize the time for all other conditions that were fit to the 2- and 3-state models. FRAP data for antibiotics in DNA or Pf phage (only one other constituent present) was fit to the 2-state model after nondimensionalizing time. The parameters found for DNA or Pf phage in the 2-state model were plugged into the 3-state model (assumes binding to be occurring independently or non-cooperatively to DNA or Pf phage) to generate the predicted FRAP recovery curves for antibiotics in DNA and Pf phage. Lastly, the DNA parameters found from fitting antibiotics in DNA to the 2-state model were kept constant, and the binding parameters for constituent 2 in the 3-state model were optimized to find the best fit to the antibiotics in DNA and Pf phage FRAP recovery curves (“Cooperative 3-state”). Numerical evaluation of each model was done in a Python Jupyter Notebook, which is available by email request.

## Statistics

All column graphs and statistical analyses were performed using GraphPad Prism version 9 software. For the bacterial killing assay, statistical significance was assessed by two-way analysis of variance (ANOVA) with Tukey’s multiple comparisons tests.

## Author contributions

Conceptualization: PLB, MJK, QC, PC, AS, EB, CM

Methodology: QC, PC, THWC, MJK, PLB, AS

Investigation: QC, PC, THWC, AG, AH

Visualization: QC, PC

Supervision: PLB, AS, CM, SCH

Writing—original draft: QC, PC

Writing—review & editing: QC, PC, EB, PLB, AS, CM, SCH

## Data and materials availability

All data are available in the main text or the supplementary materials.

## Supplementary Material

**Figure S1.**
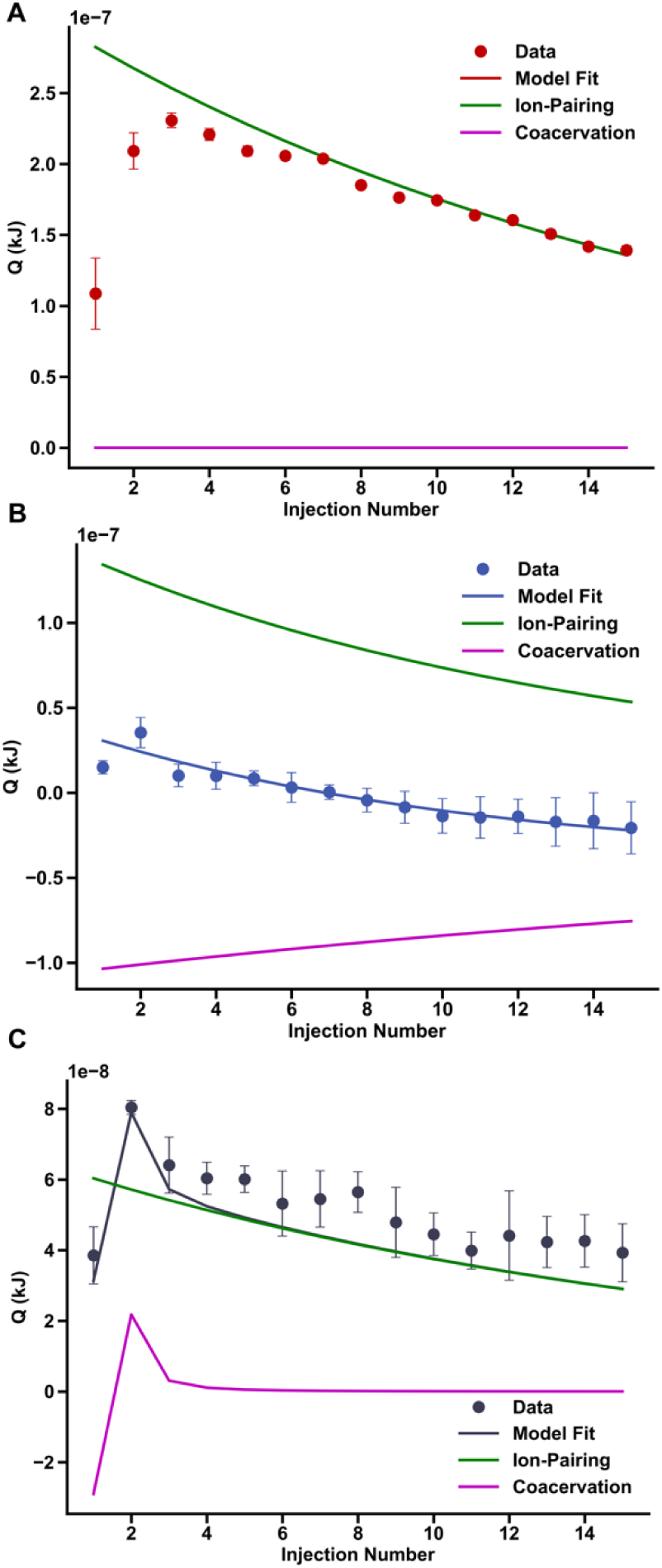
Heat released per injection from ITC measurements of tobramycin with sputum components. **(A-C)** Fitted isotherm for tobramycin in 10e11 pfu/mL Pf4, 4 mg/mL DNA, and 10e11 pfu/mL Pf4 with 4 mg/mL DNA, along with relative contributions from the coacervation and the ion pairing.

**Figure S2.**
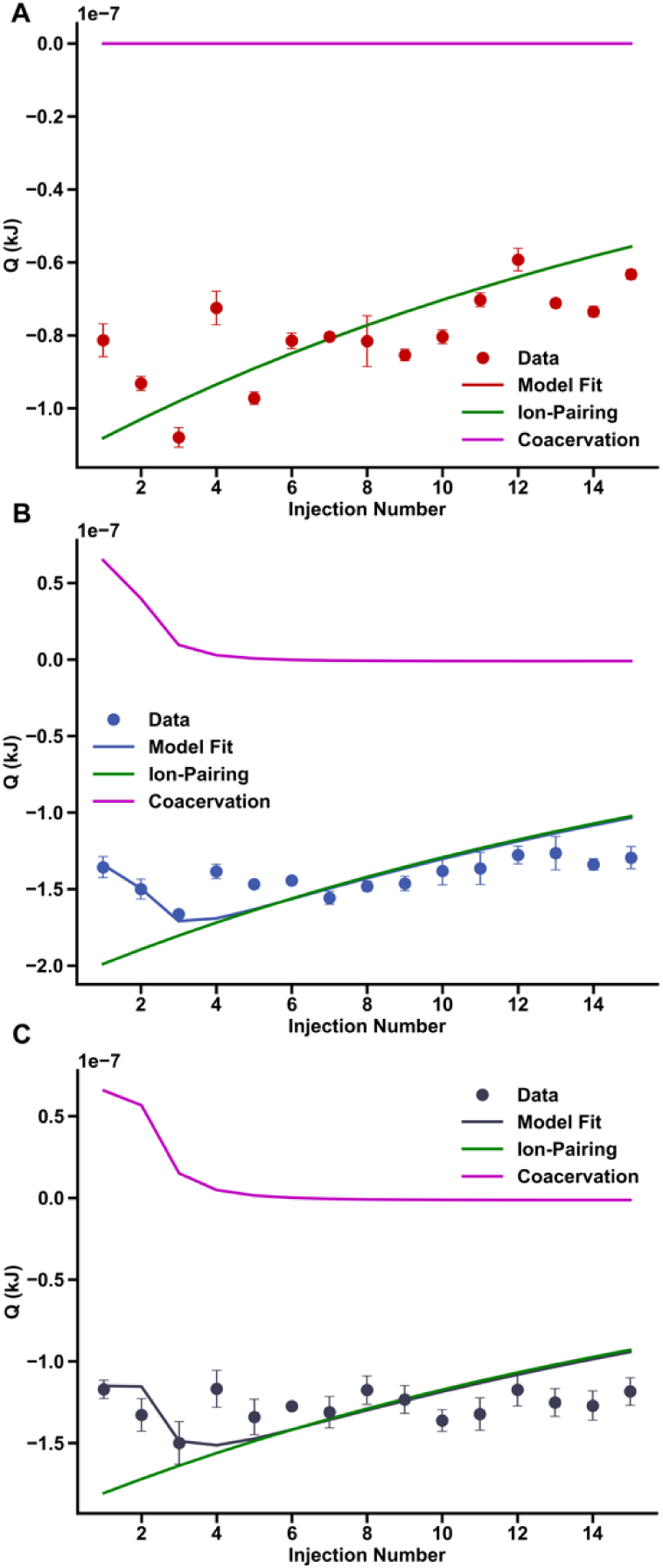
Heat released per injection from ITC measurements of ciprofloxacin with sputum components. **(A-C)** Fitted isotherm for ciprofloxacin in 10e11 pfu/mL Pf4, 4 mg/mL DNA, and 10e11 pfu/mL Pf4 with 4 mg/mL DNA, along with relative contributions from the coacervation and the ion pairing.

**Figure S3.**
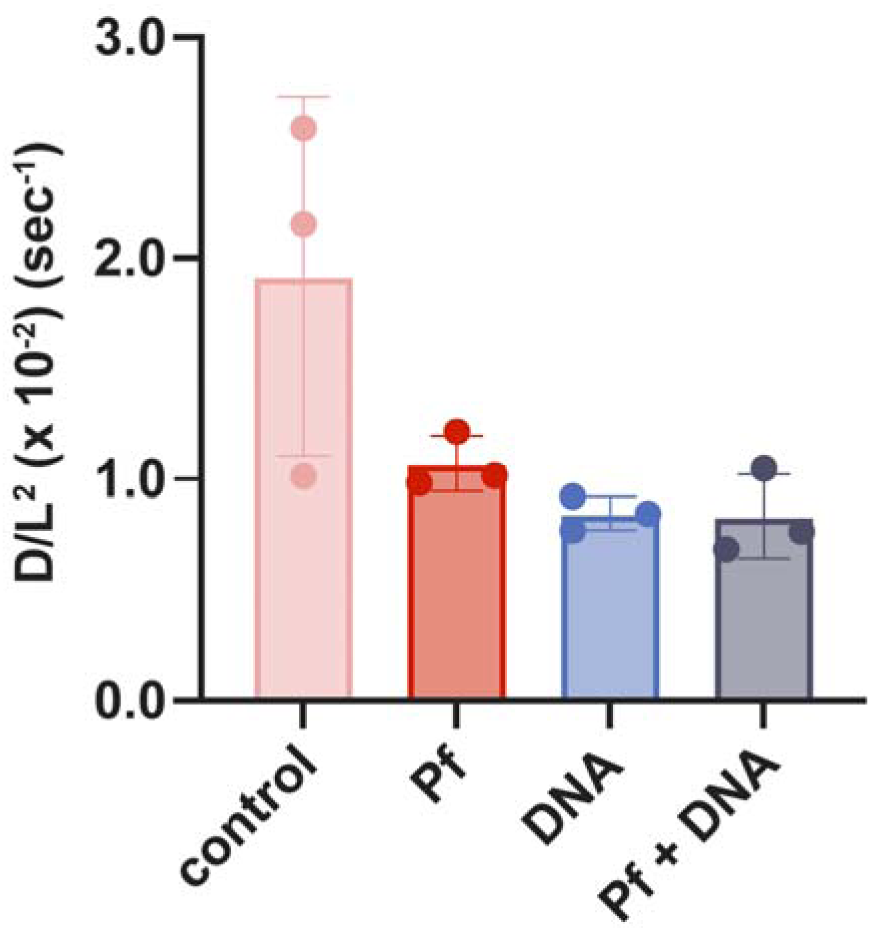
FRAP of 4kDa FITC-dextran reveals that diffusivity decreases for antibiotics are not mesh size-related (n=3). There is no statistical significance between the conditions.

**Figure S4.**
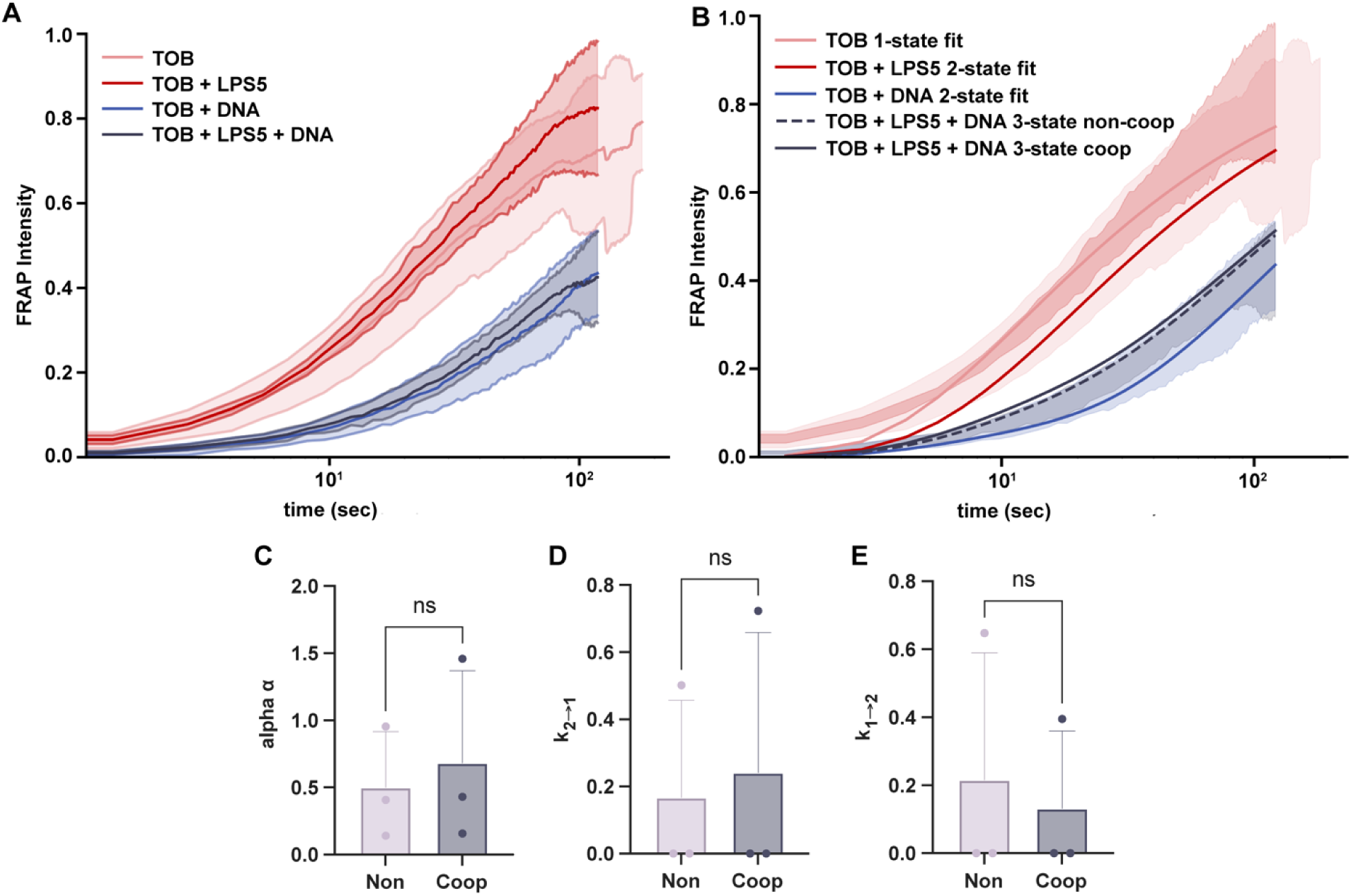
Tobramycin diffusion in non-liquid crystal-forming conditions was not affected. **(A)** Average FRAP recovery curve with standard deviation for tobramycin in non-liquid crystal forming conditions of buffer, Lytic phage LPS5, DNA, and both LPS5 and DNA. The addition of LPS5 does not significantly decrease diffusion compared to just DNA. **(B)** Fitted FRAP recovery curves for each condition with a standard deviation of data in the background. The non-cooperative FRAP 3-state model shows very little difference from the cooperative FRAP 3-state model for tobramycin mixed with LPS5 and DNA. **(C-E)** LPS5 was mixed with DNA, where the phage does not form liquid crystals with polymer. Tobramycin diffusion was not slower in solutions of LPS5 and DNA.

**Figure S5.**
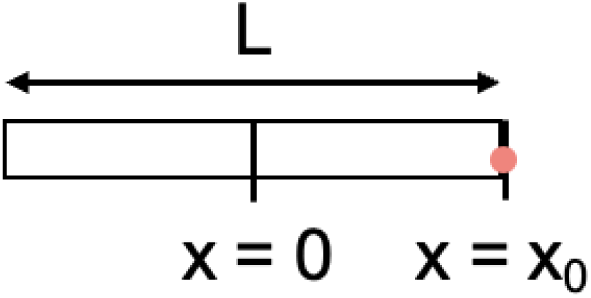
1-D line with Fluorescent Particle. A region of length L is shown with a fluorescent particle at positio x = x_0_.

**Figure S6.**
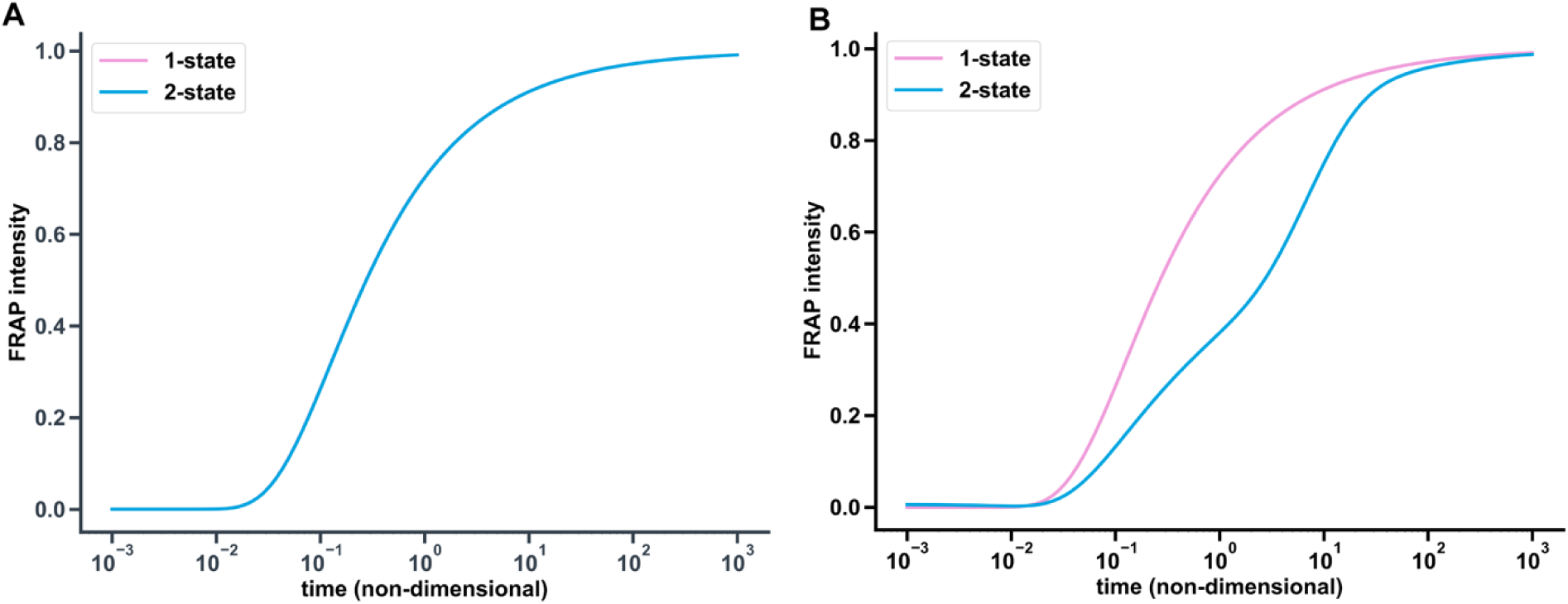
2-State Model Theoretical Output. **(A)** Setting the diffusion rate and rate constants equal for the two states in the 2-state model recovers the 1-state model FRAP curve. **(B)** When the bound state in the 2-state model experiences a lower diffusion rate, the model predicts a slower recovery curve, though eventually the curves recover to 1 with enough time.

**Figure S7.**
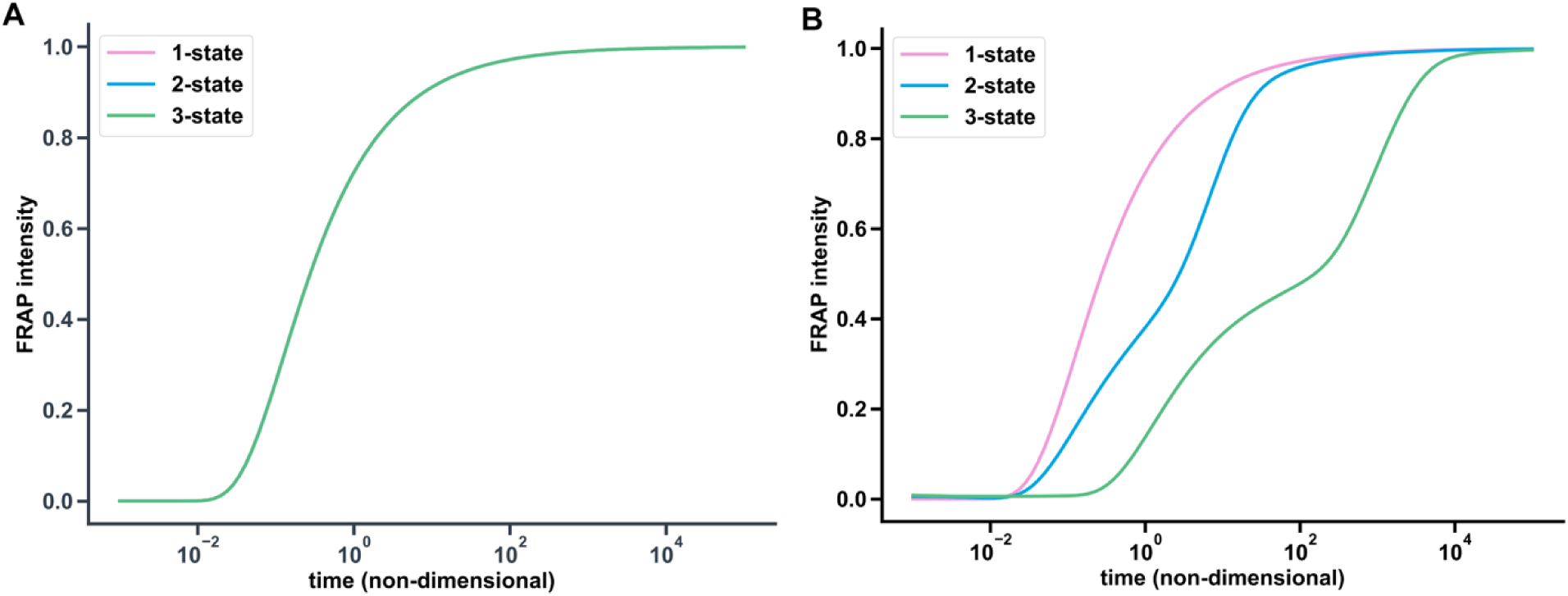
3-State Model Theoretical Output. **(A)** Setting the diffusion rate and rate constants equal for the three states in the 3-state model recovers the 1-state model FRAP curve. **(B)** When the bound states of the 3-state model experience lower diffusion rates, the model predicts a slower recovery curve than even in the 2-state model because of the additional state.

**Figure S8.**
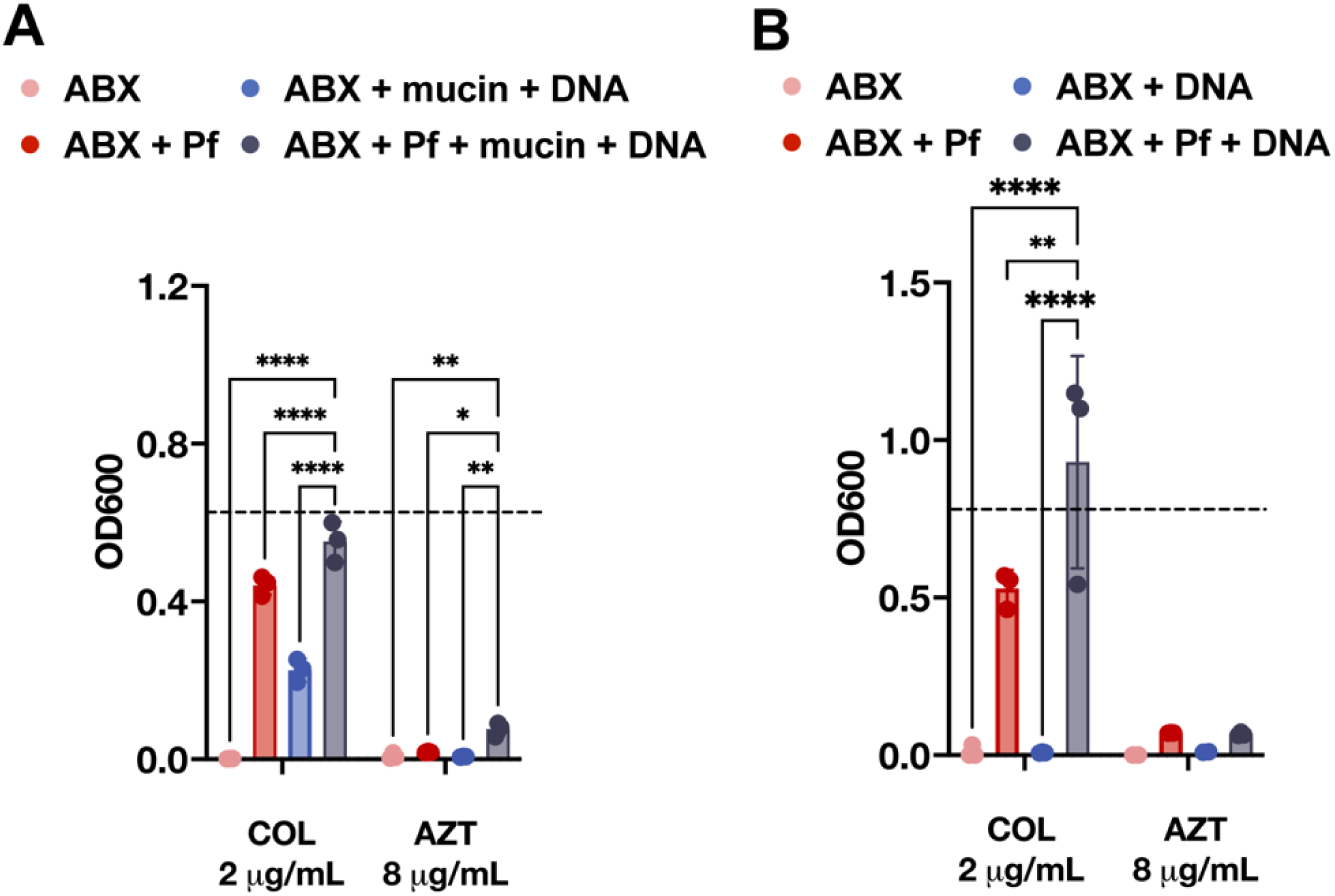
Pf and sputum polymers affect the efficacy of Colistin. To assess the impact of filamentous bacteriophage (Pf) and lung polymers of antibiotic efficacy, we measured **(A)** COL and **(B)** AZT killing of *Pseudomonas aeruginosa* (*Pa*) (n=3). antibiotics = ABX; colistin = COL; aztreonam = AZT.

**Figure S9.**
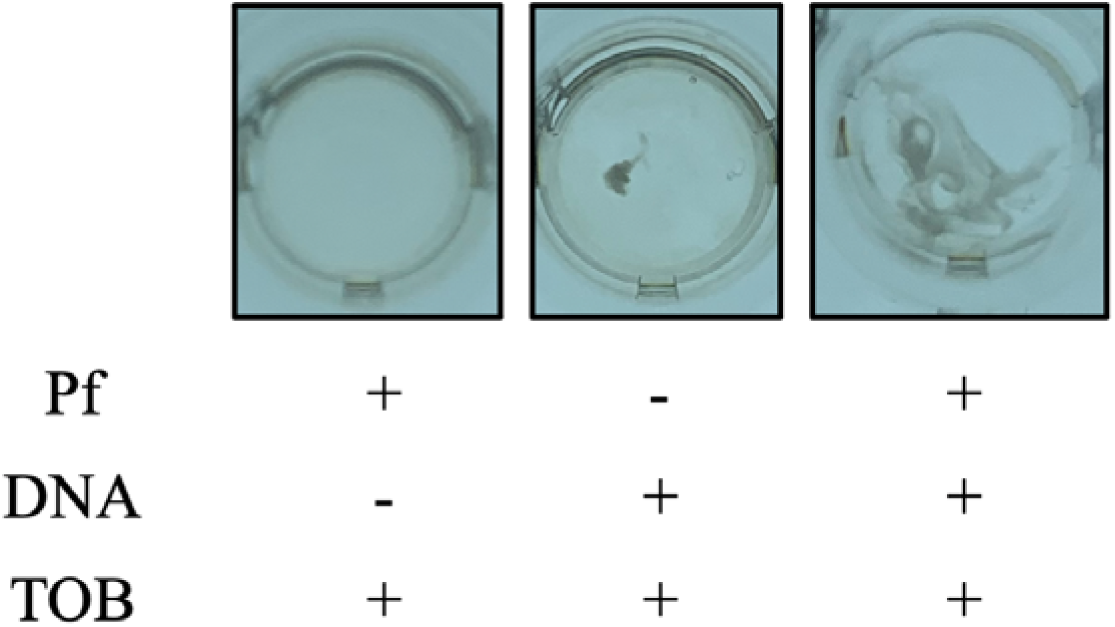
Phase change in the presence of tobramycin, Pf phages, and DNA. Pf phages (10^11^ pfu/ml), DNA (4 mg/mL), and tobramycin (1 mg/mL) were mixed as indicated above.

**Figure S10.**
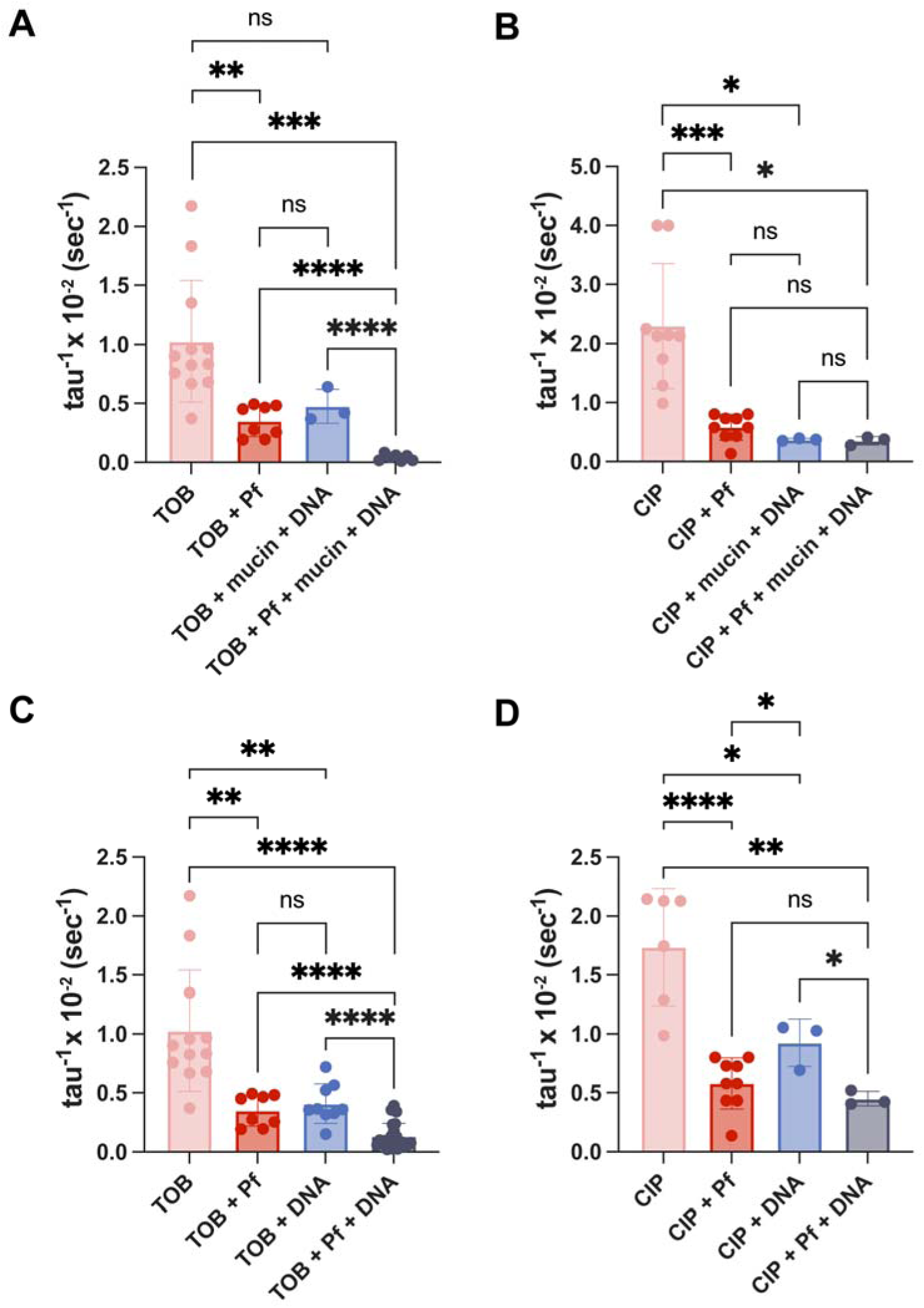
Additional pairwise statistical comparisons for effective diffusion constants derived from FRAP experiments. The effective diffusion rate, tau-1, is shown for **(A)** TOB and **(B)** CIP in artificial sputum with all comparison statistics shown (same data as **Fig. 2B, C**). The effective diffusion is also shown for **(C)** TOB and **(D)** and CIP in DNA solutions with all comparison statistics shown (same data as **Fig. 4E, F**).

**Figure S11.**
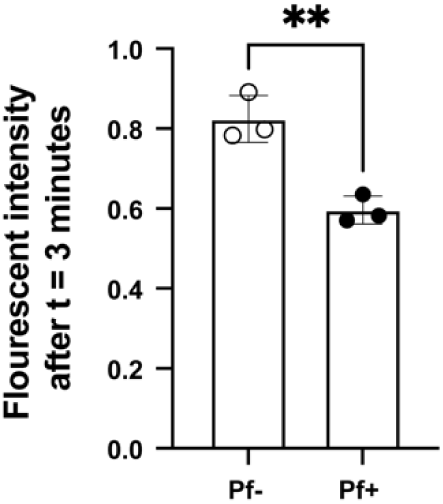
Fluorescence intensity after max recovery observation time of 3 minutes for sputum with Pf and without Pf spiked in. The fluorescence intensity of cy5-TOB after the max observation time of 3 minutes is plotted for the patient sputum samples that were and were not spiked with Pf.

**Table S1.** Patients’ information of sputum samples collected from.

| **Sputum**<br>**Sample No.** | **Gender** | **Age (years)** | ***Pa* status** | **Pf status** | **On systemic antibiotics**<br>**when sample collected** |
| --- | --- | --- | --- | --- | --- |
| **286** | Male | 31.1 | None | Negative | No |
| **334** | Female | 24.4 | Chronic | Negative | No |
| **421** | Male | 20.3 | None | Negative | No |

**Table S2.**
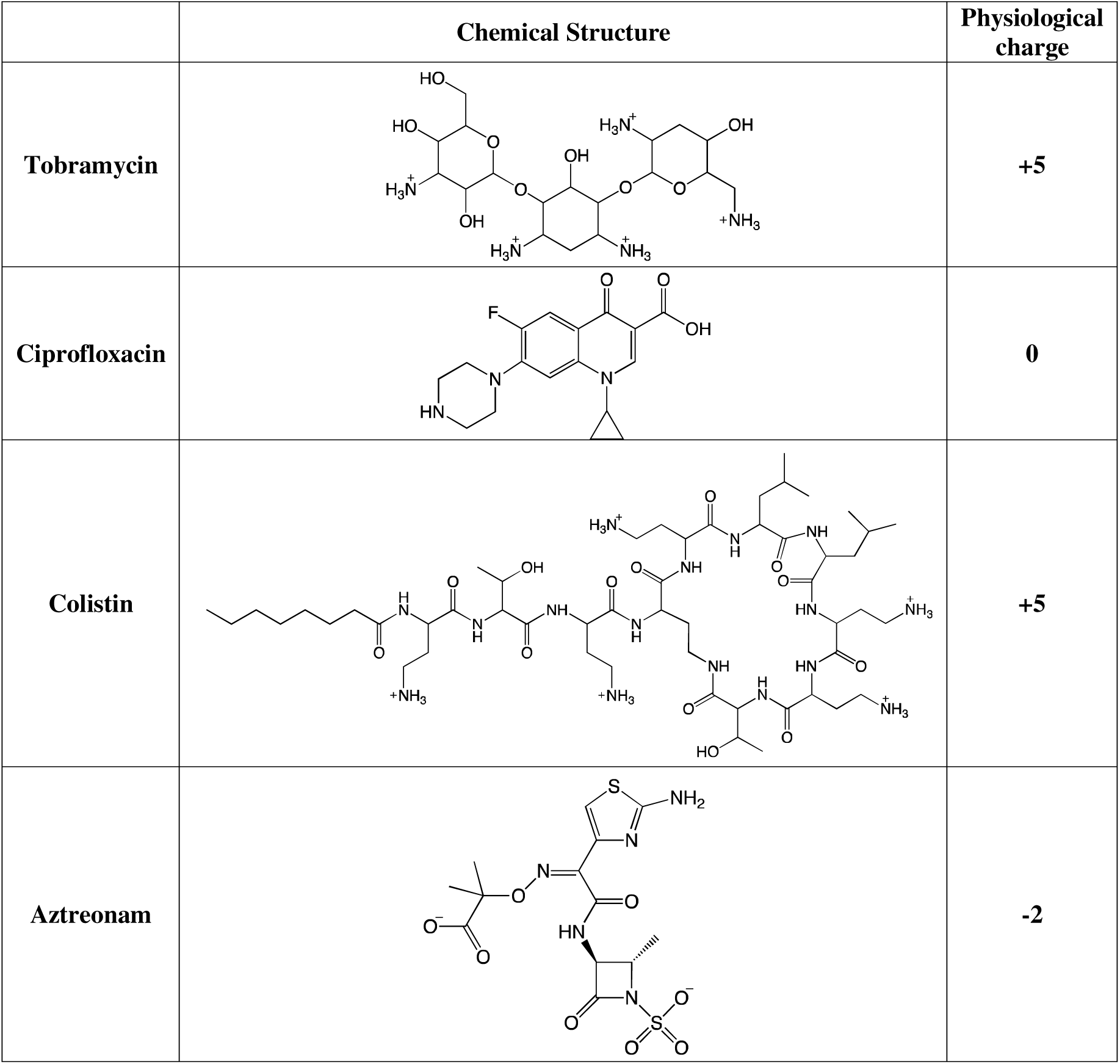
The charge of different antibiotics under physiological pH=7.

